# The geometry of knowledge in the hippocampal-prefrontal system

**DOI:** 10.64898/2026.08.19.745564

**Authors:** Manuel Schottdorf, Carlos D. Brody, David W. Tank

## Abstract

Decision making is associated with frontal brain circuits^1–5^ and spatial navigation with the hippocampus^6–8^. In addition, recent work in spatial decision making tasks^9–11^ found single neurons in both areas encoding space conjunctively with other task-relevant variables. However, circuit function is not determined by tuning alone, but also by representational geometry, i.e. the representation of task-relevant variables in neural state space. Here, using Neuropixel^12,13^ recordings in a complex spatial decision making task combined with nonlinear dimensionality reduction^8,14^, we show an intrinsically low-dimensional neural manifold in medial prefrontal cortex (mPFC) on which key task variables were represented as smooth gradients. This geometry resembled the hippocampal (HPC) map. The mPFC and HPC manifolds from one mouse can predict the behavior across other mice and brain areas. A non-linear representational map between the mPFC and HPC manifolds demonstrates alignment in time. Our work suggests that the representational geometry in HPC and mPFC is distributed and time-aligned using low-dimensional neural codes.

---

During complex behaviors, neural populations can coordinate their activity within a large neural state space onto highly organized, low-dimensional neural manifolds^15,16^. This discovery parallels a core concept in cognitive science: the internal cognitive map^6,17^. Cognitive maps organize abstract knowledge using “latent variables”. These are inferred, underlying psychological factors such as hidden task rules, the structure of an environment, or important variables that drive behavior^18,19^. Historically, latent variables in cognitive maps and internal mental spaces have been inferred from behavior alone, for example with dimension reduction^20^ or Bayesian inference^21^, suggesting that the brain can structure knowledge using geometric properties like distance, adjacency, and similarity^6,17,22^. These theoretical psychological constructs have been proposed to be instantiated in the hippocampus^7^ as non-linear, low-dimensional neural manifolds^8^. In light of the medial prefrontal cortex’s (mPFC) central role in executive function and decision-making^1–5^, we hypothesize that the neural encoding of abstract knowledge in the frontal cortex follows similar principles.

Using Neuropixel^12,13^ recordings in a complex spatial decision making task combined with nonlinear dimensionality reduction^8,14^, we discover an intrinsically low-dimensional and task-specific neural manifold in medial prefrontal cortex (mPFC) on which key task variables were represented as smooth gradients. A non-linear representational map between the mPFC and HPC demonstrated a time-aligned low-dimensional neural code, organized across both brain areas. A key factor of this alignment are neurons in one brain area, tuned to specific combinations of latent variables of the other brain area. The collective activity of these neurons forms a non-linear communication subspace. This work suggests a general principle of how distinct brain circuits organize and communicate complex information: through the temporal alignment of distributed low-dimensional and non-linear neural manifolds.

## Evidence accumulation in virtual reality

We used wildtype C57BL/6 mice (n = 8) performing an evidence-accumulation task in virtual reality (**Fig. 1A**). These mice learned to traverse the stem of an immersive virtual-reality T-maze, while visual cues were presented randomly on the left and right walls. At the end of the maze, turning to the side that had presented more cues resulted in the delivery of a liquid reward, while turning to the opposite side resulted in a time-out and an audible error signal. This ‘*accumulating towers*’ task^23^ combines navigation through a maze with decision-making, such that position and accumulated evidence (the number of left towers minus the number of right towers) both need to be represented and processed in the brain.

**Fig. 1.**
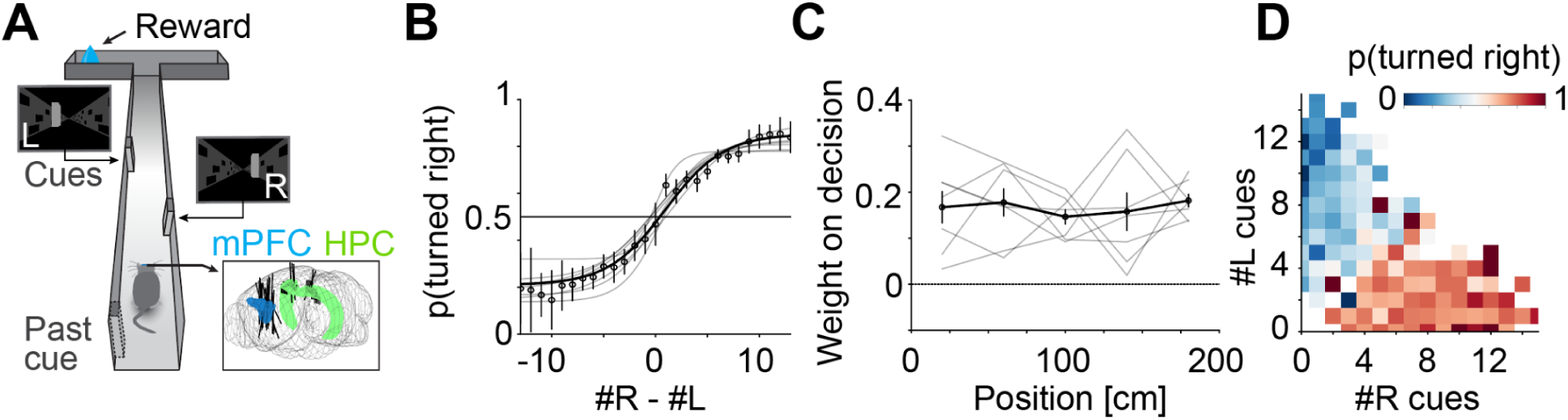
Simultaneous recordings in Hippocampus (HPC) and medial prefrontal cortex (mPFC) with Neuropixel probes in mice performing the accumulating towers task. **A)** Schematic of the task. Head-fixed mice solve a spatial accumulation-of-evidence task while neural activity is recorded with Neuropixel probes. Running along the stem of a T-maze, mice have to integrate the number of transiently visible cues, and turn in the direction with more cues for a reward. **B)** Psychometric curves of the probability to turn left relative to accumulated visual evidence, i.e. the total number of left towers minus the number of left towers. Gray lines are logistic fits to individual animals. Black dots combine data across all mice. Data are mean ± binomial confidence intervals. **C)** Logistic regression showing that mice use evidence from throughout the cue period. Gray lines are individual animals. The black line combines data across all mice. Data are mean ± s.e.m. **D)** 2D psychometrics of an example animal shows the use of both left and right cues to make a decision.

We performed acute neuropixel recordings from n = 3446 neurons (**Suppl. Fig. 1**), on average 144 ± 6 well-isolated neurons per session in mPFC and 99 ± 8 in HPC (mean ± s.e.m.). Recordings were performed simultaneously in both hemispheres and both brain areas in various combinations. The mice showed characteristic psychometric curves (**Fig. 1B**), used evidence from throughout the cue period (**Fig. 1C**) and based their decision on both left, and right cues (**Fig. 1D**).

## Joint encoding of evidence and position on a low-dimensional manifold in mPFC

Motivated by the discovery of choice selective sequences in other cortical brain areas in similar tasks^24^, we first tested if neurons in mPFC were selective for particular positions in the maze and also encoded choice. We computed the mutual information between the neural activity of each of the n=2159 well isolated mPFC cells and the position of the mouse in the maze relative to a shuffled dataset in which the activity of each cell was circularly shifted^25^. Many neurons n=794 (37%) were significantly tuned for position in the maze (see methods). Choice-specific sequences were apparent when sorting the neurons by peak position and choice of the animal (**Fig. 2A**). We next measured the mutual information between accumulated evidence and the neural activity of each cell and found that n=682 (32%) of mPFC neurons were also tuned to evidence (**Fig. 2B**). Average single cell tuning properties were generally complex (**Suppl. Fig. 2)**, and consistent with previous reports of HPC and ACC activity in this task^11^

**Fig. 2.**
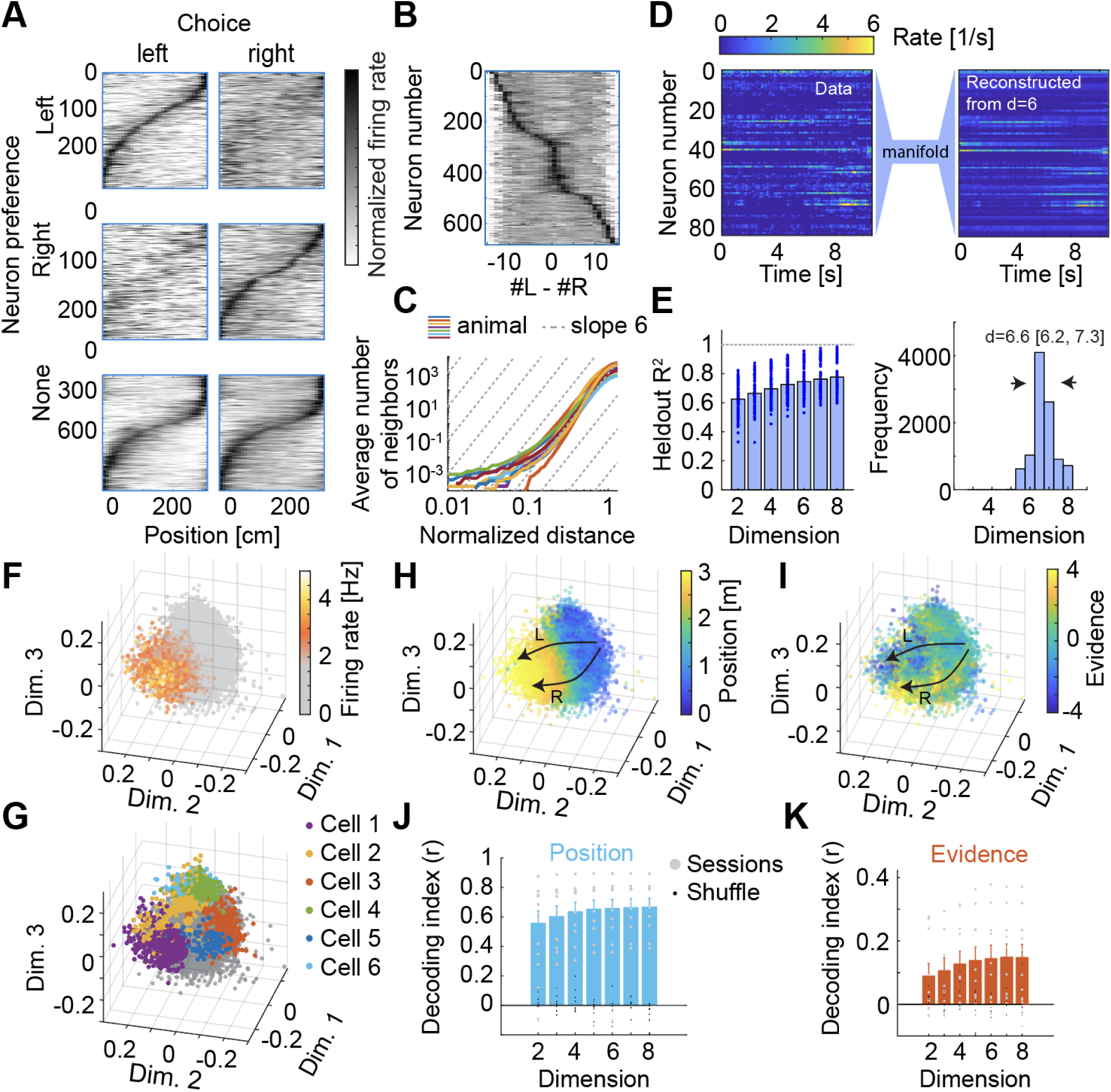
Neural activity in mPFC in the accumulating towers task. **A)** Choice-specific place cell sequences in mPFC, divided into left-choice-preferring (top), right-choice-preferring (middle) and non-preferring (bottom) cells. Cells are shown in the same order within each row group. Firing rates were z-scored for each neuron. **B**) When averaging over position, mPFC neurons also exhibit firing fields in accumulated evidence space. **C**) The mean cumulative number of neighboring neural states as a function of the geodesic distance. Grey dashed lines illustrate the slope expected for a six-dimensional manifold. **D**) Example reconstruction of held-out neural data from 6-dimensional embeddings of the neural manifold for one trial. **E**) Left: Correlation coefficient between the predicted and true firing rates in held-out trials as function of the embedding dimension. Right: Estimates of the dimensionality using bootstraps with a step-wise linear function over data shown left. **F**) Each point is a location in the three-dimensional embedding of the manifold at one time point. Coloured points represent the firing rate of one neuron. **G**) Same as F, but shown are firing rates exceeding 3σ above the mean activity for five example cells. **H)** Same as F,G, but color-coded is position in the maze. **I)** Same as F,G,H but color-coded is accumulated evidence. **J)** Decoding of position from the manifold. **K)** Same as J, but decoding accumulated evidence. Notice the curves saturating around d=6∼7 dimensions.

The measurement of tuning properties is limited to the manual selection of predetermined behavioral variables by the experimenter. We therefore turned to more principled and unsupervised methods to characterize neural activity: Neural activity can be described as a point in a high-dimensional space, in which each coordinate axis represents the activity of a single neuron. If neural trajectories are confined to a lower dimensional subregion of this space, neural activity can be described with a small number of latent variables that chart this subspace. To test this possibility for mPFC neural activity, we used manifold inference from neural dynamics (MIND)^8,14^. MIND constructs a set of latent variables with a specific emphasis on incorporating temporal dynamics and is therefore particularly suited for data with sequential activity. We first used the distance metric in MIND to estimate the intrinsic dimensionality of the neural manifold in mPFC during the accumulating towers task. We calculated distances from estimated transition probabilities between observed population activity states and counted the cumulative number of population states that fell within spheres of growing radii r, where r is an estimate for the inner distance^26,27^. If the manifold has d dimensions, we expect the number of states to grow as r^d^. We found that the number of states grows approximately as d = 5.7 (5.1–6.5; 95% bootstrapped confidence interval), indicating a low, approximately six-dimensional, latent geometry (**Fig. 2C**). To validate this estimate, we next embedded the manifold into d-dimensional Euclidean spaces and assessed how well these embedded manifolds described neural data using cross-validation on held-out trials. **Fig. 2D** shows an example of neural activity and the reconstruction of that same data from six latent variables obtained after embedding the manifold into a six-dimensional Euclidean space. Measuring the variance explained of held out data between the neural data and the reconstruction of the same data from manifolds embedded into two- to eight-dimensional Euclidean spaces, we find that the reconstruction performance saturates at around five to six dimensions. A step-wise linear fit to the various folds of the held-out data suggests a 6-7 dimensional manifold, see **Fig. 2E**. Taken together, this indicates an approximately six dimensional neural manifold produced by the circuits in the medial prefrontal cortex. This interpretation of the data makes two key predictions: First, the activity of a single neuron can be well-predicted by activity from the rest of the population and second, important task variables represented by single neurons (such as position, evidence, and choice, cf. **Fig. 2AB**) must also be represented by the population on the manifold.

To address the first prediction, we plot the activity of a typical neuron plotted as a heat map on a three-dimensional embedding of the manifold is shown in **Fig. 2F**, demonstrating a localized firing field on the manifold. Plotting the activity of multiple neurons on the same manifold reveals that many neurons tile the manifold with firing fields (**Fig. 2G**). This structure of firing fields on the manifolds implies that indeed, the activity of a single neuron can be well-predicted by activity from the rest of the population: One can infer position on the manifold from the population, and based on this population-state predict a single neuron’s activity via its manifold firing field^8^. The second prediction was the orderly representation of task variables on the manifold. **Fig. 2H** and **Fig. 2I** reveals that both position and evidence appear organized as gradients in the latent space, in that the trajectory of the neural state typically progresses along a position direction in the course of a trial, while splitting along an independent, but integrated, evidence direction. Using Gaussian process regression to decode position and evidence from the manifold, we found that both variables can be decoded from neural data, with the reconstruction score saturating around six dimensions, consistent with our earlier observations estimates of mPFC dimensionality.

## The neural manifold in hippocampus

To compare these results with neural activity in the hippocampus, we performed a similar set of analyses to our hippocampal neuropixel data. Consistent with our earlier results with Calcium imaging^8^, we found hippocampal neural activity to be characterized by choice-selective sequences (**Fig. 3A**), and also tuned to accumulated evidence (**Fig, 3B**). The number of neighbors in state space grew like a powerlaw as a function of the distance (**Fig. 3C**). Using MIND, we found that a significant fraction of neural variances is captured by few dimensions (**Fig. 3D**). Embedding HPC activity into 2-8 dimensional euclidean space, and measuring the saturation point with a stepwise linear fit across folds, suggests an approximately five dimensional neural manifold, **Fig 3E**. This low-dimensional population code is instantiated by neurons tuned to location on the manifold. An example is shown in **Fig. 3F**, and several neurons are shown in **Fig. 3G**. Similar to mPFC, we also found a smooth representation of key task variables on the manifold, see **Fig. 3H** and **Fig. 3I**, from which gaussian process regression could reliably decode position (**Fig. 3J**) and evidence (**Fig. 3K**). Note how the evidence decoding saturates at a smaller embedding dimension when compared to mPFC (cf. **Fig. 2K**), consistent with our estimate of the HPC manifold being slightly lower-dimensional than the mPFC manifold.

**Fig. 3.**
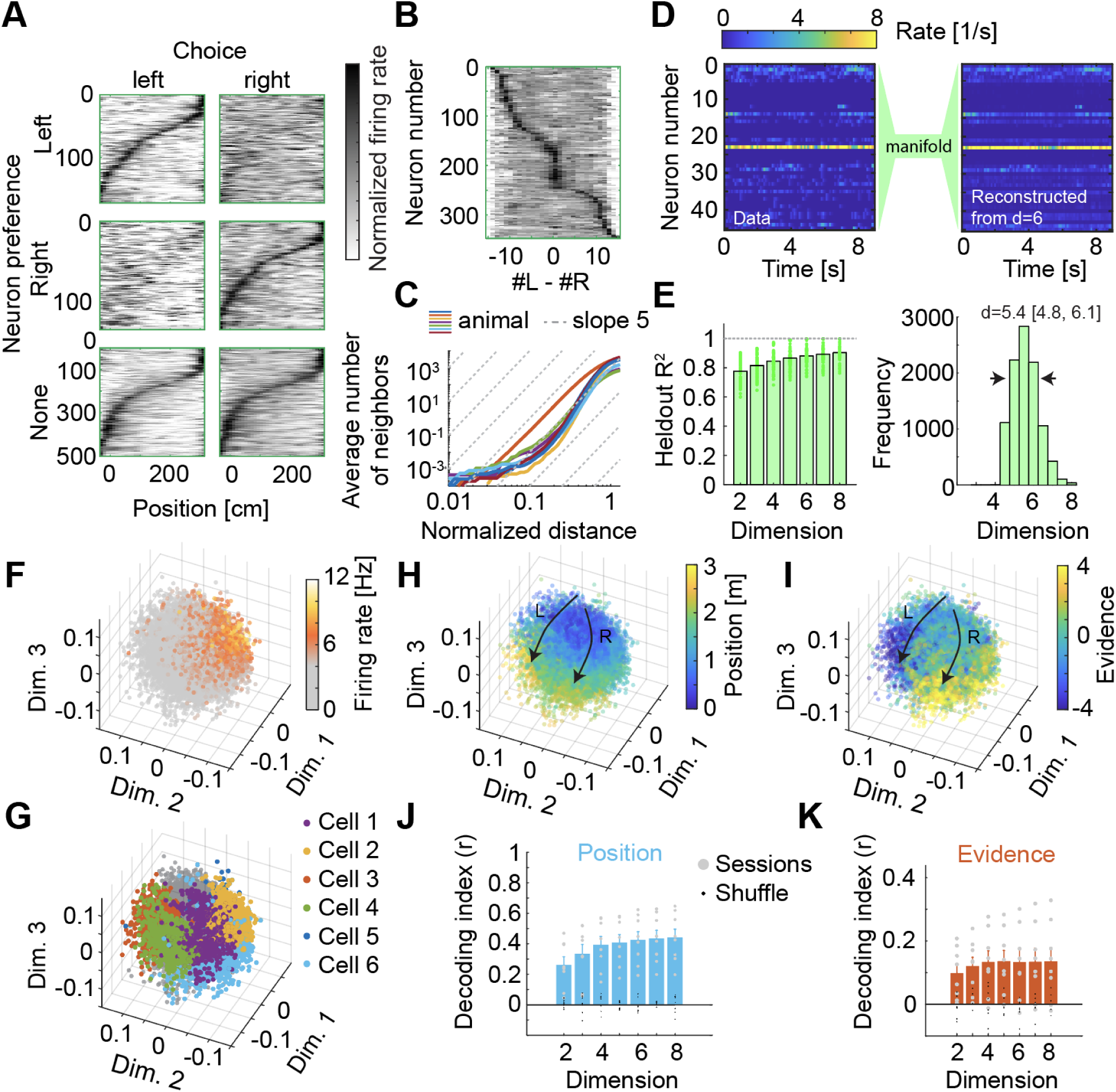
Neural activity in the hippocampus resembles mPFC and is organized on a neural manifold, **A)** Choice-specific place cell sequences in HPC, divided into left-choice-preferring (top), right-choice-preferring (middle) and non-preferring (bottom) cells. Cells are shown in the same order within each row group. Firing rates were z-scored for each neuron. **B**) When averaging over position, HPC neurons exhibit firing fields in accumulated evidence space. **C**) The mean cumulative number of neighboring neural states as a function of the geodesic distance. Grey dashed lines illustrate the slope expected for a five-dimensional manifold. **D**) Example reconstruction of held-out neural data from 5-dimensional embeddings of the neural manifold for one trial. **E**) Left: Correlation coefficient between the predicted and true firing rates in held-out trials as function of the embedding dimension. Right: Estimates of the dimensionality using bootstraps with a step-wise linear function over data shown left. **F**) Each point is a location in the three-dimensional embedding of the HPC manifold at one time point. Coloured points represent the firing rate of one neuron. **G**) Same as F, but shown are firing rates exceeding 3σ above the mean activity for five example cells. **H)** Same as F,G, but color-coded is position in the maze. **I)** Same as F,G,H but color-coded is accumulated evidence. **J)** Decoding of position from the manifold. **K)** Same as J, but decoding accumulated evidence. Notice the curves saturating around d∼5 dimensions.

## Decoding behavioral variables without behavioral measurements

Neurons in mPFC and HPC exhibit different tuning properties. While both areas give rise to choice-selective sequences, cortical tuning curves often exhibit a monotonically increasing or decreasing encoding of evidence, whereas in the hippocampus, neurons tend to have more narrow, non-monotonic tuning curves (cf. Suppl. Fig. 2 and 3; for an analysis see^11^). However, the similarity of representational geometry in HPC and mPFC suggests that the neural manifolds reflect the learned structure of the task, and are intriguingly similar across brain-areas.

To test the relationship of neural geometry to the task at hand, we hypothesize that different animals trained on the identical task must develop a similar neural geometry. To test this idea, we learn the neural manifold of animal A and, using nonlinear regression, produce a map from this latent space to a behavioural variable of interest. If the neural geometry in a different animal B is sufficiently similar, then the decoder from animal A applied to the manifold of animal B should be able to decode behavior as well.

This method is shown in **Fig. 4A**. Based on the use of similar techniques in the fMRI community^28,29^ to capture shared information across subjects by projecting neural activity into a common space, we refer to this method as Hyperalignment for short. To also test alignment across brain areas, we learn the neural manifold of brain area 1 and learn a map from this latent space to a behavioural variable of interest. If the neural geometry in a different brain area 2 is similar, then the decoder from area 1 applied to the manifold of area 2 should be able to decode behavior as well. Note that this is essentially the same technique as used above, but applied across brain areas. We refer to this method as Crossalignment for short.

**Fig. 4:**
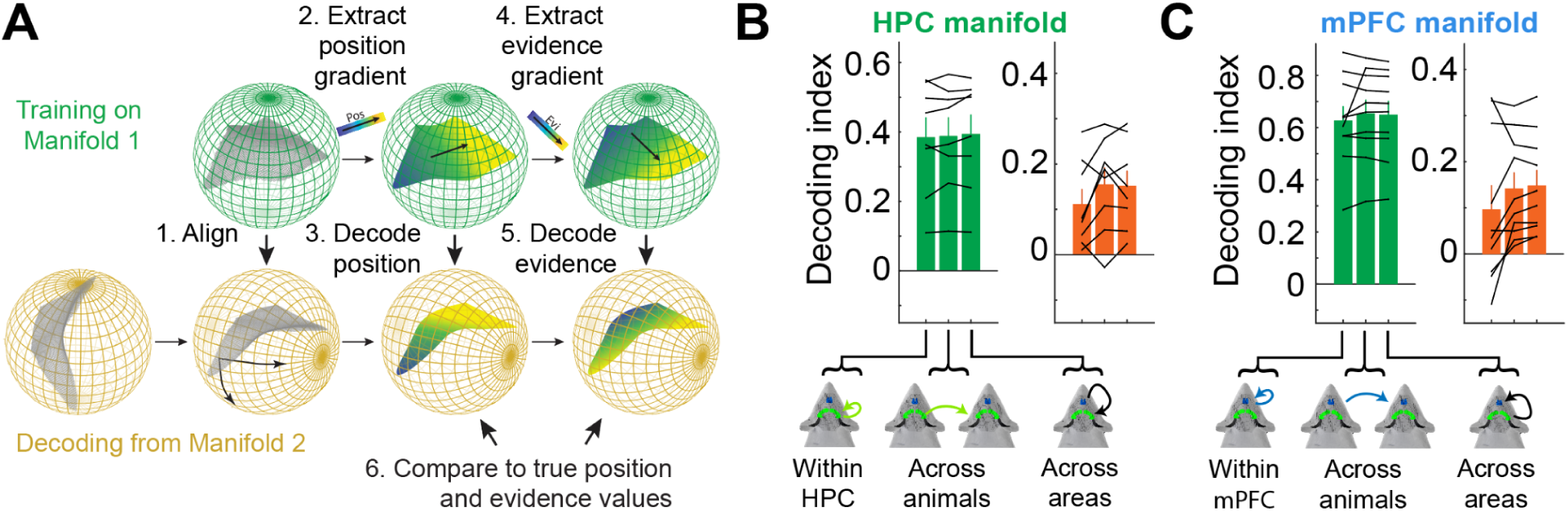
Alignment of neural manifolds across animals and brain areas. **A)** Schematic of the hyperalignment procedure: Manifold 1 (green) is used to learn a regression model of position and evidence. This model is then applied to a second manifold (orange), either from a different mouse or brain area, after optimal rotation, to predict position and evidence. **B**) Best decoding of position (left; green) and evidence (right; orange) from Hippocampus, comparing Gaussian Process Regression in the same mouse (left column) with the best hyperaligned manifold from another mouse but same brain area (middle column), and with the best crossalignment from mPFC in the same mouse (right column). **C**) Same as B, but based on the mPFC manifold.

We compared the results of these two techniques with the best decoding possible using gaussian process regression in the same animal and brain area by just splitting neural activity into training and test data. The results for the Hippocampal manifold is shown in **Fig. 4B**. We found the decoding of position (green) and evidence (orange) to be statistically indistinguishable across the three methods. This was consistent with the same analysis applied for mPFC, see **Fig. 4C**. The high degree of similarity across animals supports the hypothesis that this particular neural geometry reflects that task structure that animals learned over many training sessions. The high degree of similarity across brain areas is suggestive of a communication mechanism, in which low-dimensional neural activity produced by the resident population in area 1 allows this population of neurons to collectively interpret the coordinated activity of incoming inputs from a different brain area as movements along its own manifold. Next, we will put this hypothesis to the test.

## Precisely aligned neural geometry

The previous analyses raise the intriguing possibility that the low-dimensional manifolds in Hippocampus and medial prefrontal cortex are similar not just on average, but also time-point by time-point. To be aligned in time, two key predictions should hold true. First, for every point in time, latents charting the manifold in one brain area must be related to position on the manifold inferred from the other brain area. Second, this relationship should be strongest for small communication delays, while timing differences can suggest directionality.

**Fig. 5A-C** show the mPFC latents plotted on the HPC manifold. The latents appear highly organized. Notice the smooth gradients, resembling the encoding of behavioral variables. This alignment manifests through many mPFC neurons having firing fields on the HPC manifold: plotting the spiking of an mPFC neuron in terms of position on the HPC manifold when a spike occurred reveals such mPFC firing fields An example is shown in **Fig. 5D**, and several more neurons are shown in **Fig. 5E**. It is tempting to speculate that these across-manifold neurons can serve as anchor points for the alignment of the two manifolds. Training a decoder to predict the mPFC latents from the HPC manifold, the average best decoding saturated around 6 dimensions, suggesting that on the non-linear manifold, a majority of neural activity is shared, and private dimensions play a minor role. This alignment is only partially explained by the tuning of neurons to position and evidence in both areas. When regressing out expected neural activity based on position and evidence, and repeating the analysis, about 50% of the variance remain explained (**Suppl. Fig. 5**). The average best prediction of the mPFC latent, based on the HPC manifold for different time shifts is shown in **Fig. 5G**, and peaks close to zero. The average best prediction was found for timeshift +40 ms ± 50 ms (mean ± SEM across animals), statistically indistinguishable from zero. We also tested whether significant timing differences were present when analyzing cue and delay period separately. Across datasets, we could not find such an effect either (**Suppl. Fig. 6**). This demonstrates that the alignment is strongest for very small communication delays. This is consistent with a perfect alignment of the mPFC and HPC representation across task phases.

**Fig. 5:**
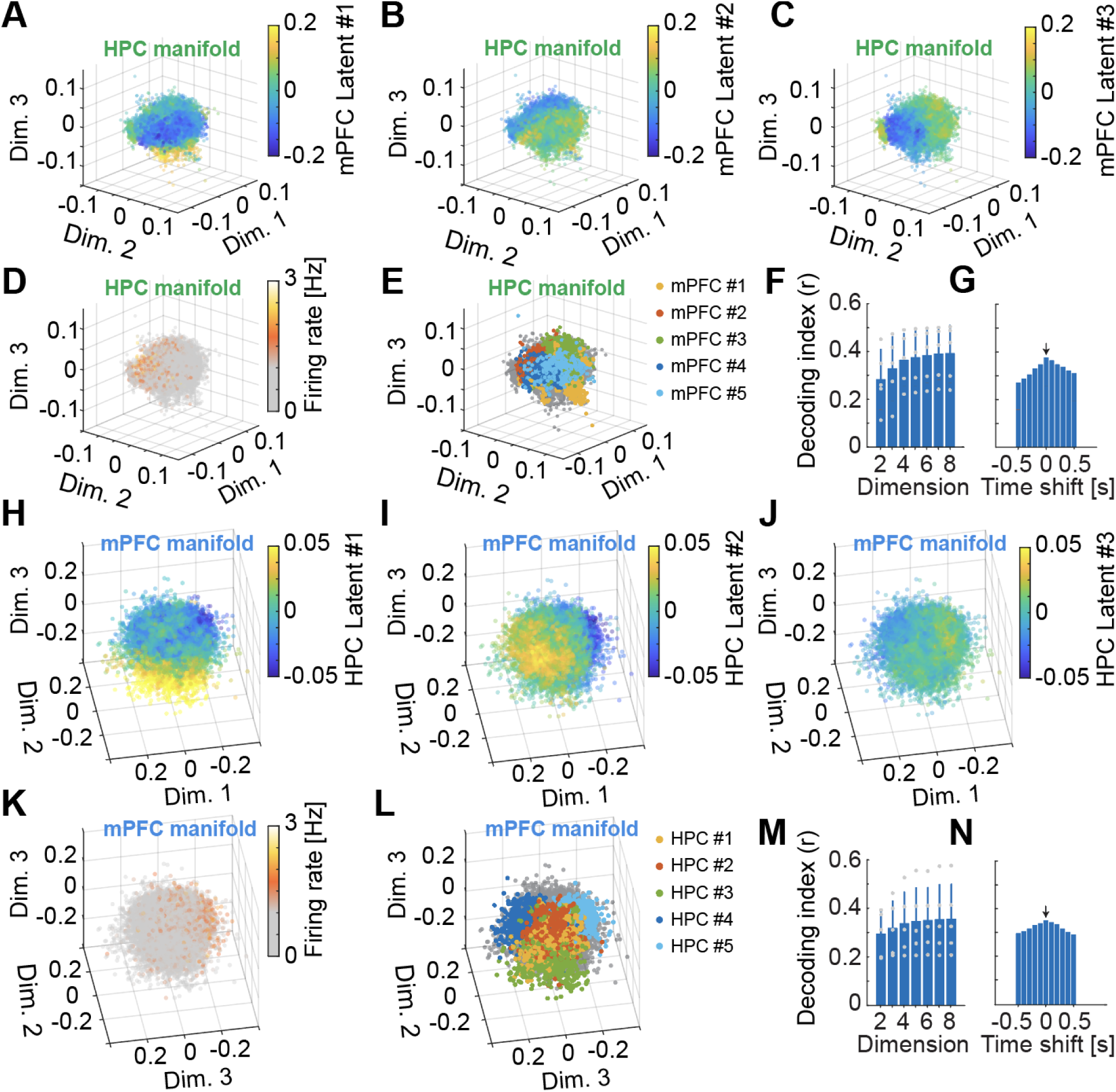
The HPC and mPFC manifolds are precisely aligned in time. **A)** Each point is a location in the three-dimensional embedding of the HPC manifold at one time point. Coloured points represent the numerical value of the first latent from the mPFC manifold. **B)** Same HPC manifold as in A, but color-coded is mPFC latent number two. **C)** Same as in A and B but for mPFC latent number three. **D)** Same HPC manifold, but color-coded is the firing rate of an example mPFC neuron. **E)** Same as D, but for five example neurons, where the activity was thresholded. **F)** Average best decoding of the mPFC latent for various embedding dimensions of the HPC manifold. **G)** Average best decoding of the mPFC latents for the d=5 dimensional HPC embedding for different time shifts. **H)** Same animal and dataset as A, but shown in the first HPC latent on the mPFC manifold. **I)** Same as H, but color-coded is the second HPC latent. **J)** Same as H and I, but color coded is the third latent. **K)** Same mPFC manifold as before, but color-coded is a single HPC neuron. **L)** Same as K but for five example neurons. **M)** Average best decoding of the HPC latents for various embedding dimensions of the mPFC manifold. **N)** Average best decoding of the HPC latents for the d=6 dimensional mPFC embedding for different time shifts.

To validate these discoveries, we performed the same set of analysis, but predicting HPC latents from the mPFC manifold. **Fig. 5G**, **Fig. 5H** and **Fig. 5I** show HPC latents on the mPFC manifold, which also form smooth gradients. Many hippocampal neurons also form across-manifold firing fields. An example is shown in **Fig. 5K**, and several other examples in **Fig. 5L**. The across-area decoding saturates at around 6 latent variables, see **Fig. 5M**, and no significant time delay was measurable. Combining both directions, we estimate that HPC leads mPFC by 10 ms ± 28 ms (mean ± SEM), again statistically indistinguishable from an exact alignment.

## Discussion

Using a virtual reality navigation task in which mice have to accumulate abstract visual evidence, we discovered that neural activity in the medial prefrontal cortex, on the single-trial level, is organized as a low-dimensional and task-specific neural manifold. On this nonlinear manifold, individual mPFC neurons formed firing fields and the key task variables of accumulated visual evidence and spatial position were arranged orderly (Fig. 2). This was accompanied by an even lower-dimensional manifold in the hippocampus (Fig. 3). These representational structures were remarkably similar across animals (Fig. 4) and aligned in time (Fig. 5). Previous studies^9–11^ have shown that trial-averaged tuning curves of single neurons in both areas can encode space conjunctively with other task-relevant variables. In our task, trial-averaged individual neurons in HPC and PFC exhibit different evidence tuning^11^. Yet, we discovered a remarkably similar geometry of the MIND-inferred manifolds on the single trials level; stronger than what would be expected from position and evidence tuning alone. This suggests a tight coordination across neurons and across regions beyond the known experimental variables of position and evidence on a trial-by-trial basis. This alignment could be computationally beneficial. Circuits in the hippocampus and frontal cortex were suggested to perform the accumulation of evidence independently with different neural circuits^11^, serve cognition with different mnemonic algorithms^30,31^, contain information in an abstract format^22^, or operate on different time scales^32^. Our empirical findings might provide a mechanism by which these potentially different computational roles are integrated to serve behavior.

Distributed geometric representations of learned knowledge, produced by sequential neural activity of neurons with highly organized firing discharges across brain areas, could serve as an organizing principle of how brain areas communicate and integrate results of local computations to meet task demands. It is tempting to speculate that these mechanisms that organize learned knowledge can play an important part in bridging the gap between microcircuits and the cognitive science of whole-brain networks^33^. Our work also suggests that the preserved neural dynamics across animals performing similar behaviour^15^ could possibly generalize across brain areas, using the same “anchor neurons” to coordinate cognitive processes. Finally, recent work demonstrated Hippocampal-Retrosoplenial communication in similarly organized subspaces^34^ and offline replay demonstrated temporally organized sequences of cortico-hippocampal interactions^35,36^. Intriguing parallels exist between these communication subspaces, replay events and the latent variable structures discovered here. Future work, particularly during learning, will be key for a quantitative understanding of how these representations form and align.

## Acknowledgements

We thank Alvaro Luna and Christian Tabedzki for maintaining data infrastructure. Lindsey S. Brown, Mika E. Diamati, Joshua B. Julian and Jesse Kaminsky provided very valuable feedback. We also thank the Tank and Brody labs for discussions. Laura A. Lynch, Sanjeev R. Janarthanan, and Samuel S.-H. Wang maintain the PNI imaging core where the light sheet imaging was performed. This work was supported by NIH grant U19NS132720, as well as the Simons Collaboration on the Global Brain. MS is supported by a CV Starr fellowship, and a Burroughs-Wellcome Fund Career Award at the Scientific Interface (CASI). M.S., C.D.B. and D.W.T. designed the experiments. M.S. designed and built the rig, performed the experiments, analyzed the data and wrote the manuscript. M.S. C.D.B. and D.W.T. revised the paper. C.D.B. and D.W.T. supervised the project.

## Methods

### Experiment and data processing

#### Animals and stereotaxic surgery

All procedures performed in this study were approved by the Institutional Animal Care and Use Committee at Princeton University under protocol 1910 and were performed in accordance with the Guide for the Care and Use of Laboratory Animals. n=3 male and n=4 female C57BL/6 mice were used in total. During training and recording, their ages ranged between 3.5-10.5 months; median 6.4 months. Mice were housed under a 12-hr/12-hr light/dark cycle. All experiments and surgical procedures were conducted during the dark cycle.

Surgeries were performed under aseptic conditions and body temperature was maintained with a heating pad (Harvard Apparatus). Mice were anesthetized with isoflurane (3% for induction, 1–1.5% for maintenance) and given a preoperative dose of meloxicam subcutaneously for analgesia (10 mg/kg) and a postoperative dose 24 h later. After asepsis, the skull was exposed, and the periosteum was removed. Four small dents were drilled into the skull for use as fiducial marks. These were made with a pneumatic drill over CA1 (mediolateral, ±1.8 mm from the midline; anteroposterior, 2.0 mm posterior from bregma) and mPFC (mediolateral, ±0.4 mm from the midline; anteroposterior, 1.8 mm anterior from bregma). Then a gold coated ground pin (Newark 82K7797) was affixed above the cerebellum, approximately 6.0 mm posterior and 2.0 mm lateral from Bregma on the left side. Finally, a custom lightweight titanium head-plate (∼1g weight) with an opening in the center section was attached to the skull with a transparent adhesive cement (C&B Metabond, Parkell). Transparent Metabond was chosen to not obfuscate the fiducial marks.

Mice were allowed to recover for at least 5 days before starting water restriction for behavioral training. Mice were extensively handled during the restriction process to familiarize them to experimenters. Mice were allotted daily volumes of 1–2 ml of liquid per day, delivered either during behavioral sessions or supplemented after sessions. Mice were examined daily to ensure that there were no signs of dehydration and that a body mass of at least 80% of the initial value was maintained. Mouse husbandry information, body weight, and water reward history was automatically ingested and stored in a relational database, interfaced through datajoint^37^. All training was performed in custom VR systems (see **Suppl. Fig. 1A)**.

A second, very brief, surgery was performed after training when mice had reached a performance level of >60% correct. Mice were again anesthetized with isoflurane (3% for induction, 1–1.5% for maintenance) and given a preoperative dose of meloxicam subcutaneously for analgesia (10 mg/kg). After asepsis, small craniotomies (<1mm) were drilled centered around the fiducial marks. Care was taken to not damage the dura in the process. The craniotomies were then sealed with a bio-inert silicone polymer (Kwik-Sil) and the animals were given 24h to recover in their home cage. Mice recover very quickly (within minutes) and are not taken off water restriction during the recovery period. After a recovery period of at least 24h, daily recordings were made. To this end, the mouse is head-fixed in the apparatus, probes are inserted under visual control through a stereo microscope with long working distance (see **Suppl. Fig. 1B**). Details of the recordings are provided below.

After no more than one week of acute recordings, animals were perfused transcardially and their brains cleared using iDisco. Mice were deeply anesthetized (200 mg/kg ketamine and 20 mg/kg Xylazine) and then transcardially perfused with ∼100ml of phosphate buffered saline (PBS) supplemented with 20 units/ml Heparin followed by ∼100ml of 4% paraformaldehyde (PFA) in PBS. Brains were removed from the skull and post-fixed for 24h in 4% PFA at room temperature. They were then washed several times and stored in PBS for iDisco clearing and light sheet imaging.

#### Behavioral training

25 mice were trained to perform the accumulating towers task in a virtual-reality environment, as previously described.^8,23^ In brief, mice were headfixed so that they could run comfortably on an 8-inch (20 cm) Styrofoam ball suspended by an air flow of ∼70 liters per minute between the ball and a custom 3D printed cup. Ball movements were monitored with an optical flow sensor (ADNS3080) in the cup, connected to an Arduino Due. The virtual-reality environment was projected onto a paint-coated Styrofoam screen (approximately 270° horizontal and 80° vertical visual field) using a consumer-grade projector (Optoma HD28HDR). In contrast to our imaging studies^8,38^, we did not employ color-filtering in the projection system. The mice experienced the VR world as gray. A CAD rendering of the system is shown in **Suppl. Fig. 1A**. The light produced by the Optoma projector had significant overlap with the animals’ visual sensitivity (**Suppl. Fig. 1C**). The virtual environment was generated using the ViRMEn software^39^. Rewards were delivered by a solenoid valve (NResearch), controlled by a NI USB-6501 (National Instruments). Behavioral information was recorded in files, also automatically ingested into the same database mentioned above interfaced through datajoint^37^.

Mice were trained to run down a 330-cm virtual T-maze (30-cm start region, 200-cm cue region and 100-cm delay region). As mice ran through the cue region, tall, high-contrast visual cues (towers, 6 cm tall and 2 cm wide) were shown along either wall. After the delay period, mice were presented with a liquid reward for turning into the arm on the side where more towers had been shown (4–8 μl of 10% w/v sucrose). Rewarded trials were followed by a 3-s inter-trial interval, and error trials were followed by an audio error cue and a 12-s inter-trial interval (ITI). When rewards or error cues were delivered, the visual display froze for the first second after which the display was then blacked out. Tower positions were drawn randomly from spatial Poisson processes with means of 7.7 and 2.3 towers per meter on the rewarded and unrewarded sides, respectively. Towers were transient, appearing when mice were 10 cm away from their locations and disappeared after 200 ms. Each session started with at least 10 trials of a visually guided version of the task as warm-up before proceeding to the main task. On the final level the median duration of a session was 61 min (60min - 78min for 68% range; n=7 mice). The median trial duration for the seven mice was 11.6s (9.3s - 18.9s for 68% range, including ITIs and timeouts for error-trials). The median duration of the cue period was 2.2s (1.8s - 3.8s for 68% range) and 2.0s for the memory period (1.3s - 3.0s for 68% range). Detailed methods for the shaping procedures involved in training mice to perform the task, as well as performance and behavioral analyses, have previously been published^23^.

Out of the 25 mice that entered the training pipeline, 10 mastered the accumulating towers task in six weeks (which was chosen as maximum time of training). In all 10 mice, acute recordings were performed in various combinations of brain areas. Out of these animals, 7 mice produced good behavior in the 7-day acute recording window, and electrodes were successfully placed in mPFC and HPC. The final seven animals included in this study performed 3754 daily behavioral sessions in total, including the shaping procedure and sessions with neurophysiological data collection.

#### Acute neuropixel recordings

The mouse was head-fixed on a custom-built VR recording rig. Then the Kwik-Sil was removed and the craniotomy cleaned with sterile saline. CM-DiI (Invitrogen C7000) coated Neuropixel probes were angled at ∼10 deg along the ML axis for the Hippocampus, and ∼10 deg along the AP axis for mPFC. To maximize neuron count in mPFC, we use two NPX1.0 probes^13^ with 384 active sites each to record from both hemispheres. Two NPX2.0 probes^12^ (test-phase, four-shank, Imec) were used for simultaneous HPC recordings in both hemispheres with eight shanks positioned in HPC. The probes’ ground and external references were connected through a silver wire that terminated in a gold pin mating with the pin implanted above the cerebellum of the mouse. The probes were then manually lowered through the craniotomies to just above the surface of the brain under visual control through a stereo microscope (Zeiss Discovery V.8; 0.3x objective with 23.6mm working distance). The probes were then covered with a drop of sterile saline (0.9% NaCl) and a layer of silicone oil to prevent drying. The probes were then automatically inserted (10 µm/sec) with a set of four 3-axis manipulators (New Scale Multi-Probe Micromanipulator) that were integrated into the VR rig (see **Suppl. Fig. 1**). For HPC recordings with Neuropixels 2.0 (active length 720 µm), probes were inserted 1.9 mm, and retracted 100 µm. For mPFC recordings with Neuropixels 1.0 (active length 3.84mm), probes were inserted 4.1mm, and retracted 100µm. We recorded 384 channels per probe. For NPX1.0 probes, we chose the channels closest to the tip of the probe. For 4-shank 2.0 probes, we selected the 96 channels closest to the tip across the four shanks. The insertion of four probes takes ∼30min, after which we allowed the preparation to stabilize for 15 min before starting the behavior session.

The VR emitted a binary pattern of TTL pulses (NI USB-6501) that encoded the frame number. These pulses were recorded in SpikeGLX using an auxiliary National Instruments data acquisition card (NI PXIe-6341 with NI BNC-2110) to synchronize VR traces with neurophysiological data. This card and the Imec card were placed in the same PXIe-1071 chassis.

After each recording session, the probe was rinsed with MilliQ water, soaked in an enzymatic detergent (fresh 1% Tergazyme in MilliQ water) overnight at room temperature, then rinsed and soaked again in MilliQ water for at least 15 minutes before dry storage. Before recordings, probe tips were coated with CM-DiI (Invitrogen C7000). The dye was dissolved in IPA to 1 µg/µl in IPA, a drop placed on the probe tip under a dissection microscope (Leica Stereozoom S9E), and left to dry.

#### Data processing

Spike sorting, synchronization and data processing were done as follows. Neuropixel data was acquired at 30 kHz. For NPX1.0 probes we used the 250/500 LFP/AP gain setting. VR data were synchronized to spiking data by computing the current VR frame number from the pattern of TTL pulses recorded simultaneously to the NPX data. The drift between the NI PXIe-6341 and the IMEC PXIe over the course of a full experimental session was reduced to ∼50ms by precise measurements of the respective clocks. The residual error is similar to the delay of the video projector, used in the VR system itself. Neuropixel data was preprocessed with CatGT to apply global common average referencing (-gblcar) and to isolate the action potential frequency band (-apfilter=butter,12,300,9000). Spike sorting was then performed offline using Kilosort. This pipeline was implemented in datajoint and ran automatically over newly generated files: New files produced by SpikeGLX were automatically copied to an in-house cluster using Globus. On this cluster, equipped with GPUs, a set of slurm scripts automatically ran CatGT, Kilosort, and the custom synchronization code. The relevant VR information was automatically queried from the database using datajoint. When sorting and synchronization was complete, the results were stored in the database. Finally, at a local workstation, we used Phy to remove clusters with obvious irregularities such as noise and classify clusters as single-unit (“good”). All unit isolation and manual curation was performed blind to the session’s corresponding behavioral data. After inspection, the spike trains were first causally smoothed with exponential kernels of width τ = 100 ms, and then resampled at 100 ms bins. The resulting firing rates were exported from python and stored as .mat files for further processing.

#### Histology and probe tracking

PFA-fixed brains were cleared with iDISCO+ and used to acquire high-resolution and high-signal-to-noise whole-brain light sheet imaging volumes. Imaging data in the 488 nm channel was used for autofluorescence, and 561 nm for CM-DiI. The 488 channel was used to align the imaged volume to the Allen atlas. Fluorescence at 561 nm was used to infer electrode tracks. Histology was used primarily for the mPFC electrodes. For placement in Hippocampus, theta power in the LFP band proved highly informative. Histology was used as additional confirmation of correct electrode placement. Neurons positioned in Anterior cingulate, Prelimbic and Infralimbic were pooled for mPFC activity. HPC data includes CA1 and Dentate (cf. **Suppl. Fig. 4**).

### Analyses

#### 1D Psychometric curves

Psychometric curves (Fig. 1b) were plotted using previously described methods. In brief, psychometric curves were fitted using a four-parameter sigmoid: p(Δ) = b + a/(1+exp(-(Δ-d)/l)) in which Δ is the difference between the number of right and left towers. The parameters “a” and “b” capture the sensitivity to the towers. “d” captures a bias, and “l” the width of the psychometric curve. The fits were done with curve_fit from scipy.optimize on the mean behavioral data, binned into discrete evidence bins of size 1. Errorbars in Fig. 1b show two times the standard error of the mean, approximating 95% confidence intervals.

#### Logistic regression analysis

Logistic regression was performed using previously described methods. In brief, we modelled choices of the mice in each trial with logistic regression in which the factors are the evidence (number of right towers minus number of left towers) in five equally sized regions in the cue period. The fit was done with LogisticRegression from the sklearn.linear_model package. Data points and error bars in Fig. 1c show mean and standard error of the mean across the N=7 mice. Thin gray lines are the individual animals. As these behavioral analyses require considerable amounts of data, Fig. 1b and Fig. 1c includes the training data as well.

#### 2D Psychometrics

For 2D psychometrics (Fig. 1c), we show data from an example animal (M022, female, all data from T11). The animal’s choices were displayed in a 2D space with the number of left cures, and the number of right cues on both axes. Every bin shows the fraction of trials in which the mouse turned right. Other animal’s psychometrics are similar. To demonstrate that animal behavior is unbiased on a single-animal level, we display a single example animal in Fig. 1c.

#### Mutual information analysis

To produce sequence plots (Fig 2A,B and Fig. 3A,B), similar to our earlier work, we evaluated for each cell a previously defined mutual information metric^25^, I = ∫ λ(x) log_2_(λ(x) / λ) p(x) d^2^x. I is the mutual information rate of the cell in bits per second, x is the spatial location of the mouse, λ(x) is the mean firing rate of the cell at location x, p(x) is the probability density of the mouse occupying location x. λ(x) was then calculated bin-wise by collecting all smoothed firing rate values in their respective bins across the entire session and taking the mean. p(x) was calculated similarly by counting the number of frames that the mouse spent in each bin across trials and normalized to have a sum of 1. The mean firing rate for each cell was then computed as λ = ∫ λ(x) p(x) d^2^x

For position data, 10-cm bins were used. For evidence data, 31 bins (–15 to 15, number of right towers minus number of left towers) were used. To determine significance, the mutual information value of each cell was compared with the mean mutual information value of a shuffled dataset (100 shuffles), in which the firing rates of each cell was circularly shifted by a random interval within each trial, which disrupts the relationship between position and neural activity, but maintains neural activity patterns. Only cells that had mutual information values greater than 2σ above the average mutual information of the shuffle distribution were considered statistically significant. Cells with statistically significant mutual information between neural activity and position in left-choice trials, but not right-choice trials were categorized as left-choice preferring, whereas cells with statistically significant mutual information between neural activity and position in right-choice trials, but not left-choice trials were categorized as right-choice preferring. Those that were significant for both left and right choice trials were categorized as non-preferring. The same analysis was done for mutual information between neural activity and evidence. To obtain sequence plots, firing rates were z-scored, and ordered based on the position of the peak mean firing rate. Trials that entered into the analysis were selected based on block performance only. We included all blocks in which the average performance exceeded 60% correct.

#### Manifold inference from neural dynamics

Consistent with the previous analysis, we restricted our analysis to neural data in cue and delay periods in blocks exceeding 60% correct trials and to neurons of which the firing rate between first and last third of the recording did not change by more than 75%. This (mild) thresholding eliminates the few cases of units close to the spike sorting threshold that were lost over the course of the session. We then followed the previously published procedure to calculate the distances between pairs of population activity vectors, extracting a set of latent variables from these distances with multidimensional scaling, and learning a map between latent space and network activity with local linear embedding (LLE).^8^

In brief, we first learned a generative model of transition probabilities from population firing rates s(t) = [s_1_(t), …, s_N_(t)] of N neurons at time 0 < t < T, to the activity s(t + Δt) using the previously developed random forest method^14^. When splitting the neural state space into regions using a set of hyperplanes organized in a decision tree, we assessed 20 random hyperplane orientations at every node of the tree and selected the orientation that best split the data and we set the minimum number of leaves in each random tree to 1000. To define transitions, we considered all states Δt = 100 ms apart. Finally, to make the manifold fitting computationally tracktable, we fit the manifold only to 50% of the data identified with the previously published landmark algorithm^14^. All other hyperparameters were chosen as previously described. The random forest model provides us with a set of transition probabilities p(s(t + Δt)|s(t)) that can be translated into a local distance δ(s(t + Δt), s(t)) under a diffusion approximation, in which the transition probability p decreases with distance δ as p ∝ exp(−δ^2^). Similar to isomap, we then calculated the global distance between two states as the length of the shortest path from one to the other via any intermediate, connected states. The pairwise geodesic distances of l points ρ(i,j), in which 0 < i, j ≤ l, then yields a matrix of size l × l that was embedded using multidimensional scaling with Sammon’s nonlinear mapping. This yielded latent variables to describe population data. The mapping from latent space to neural activity and back was then achieved with local linear embedding.

The MIND algorithm is computationally expressive, and can lead to overfitting. To ensure reliability of the fits, we used cross-validation, in which the manifold was fit to all data, except a set of held-out trials. These held-out trials were then used to measure the performance of mind (cf. Fig. 2D,E and Fig. 3D,E). The same analyses and parameters were used for mPFC and HPC neural data.

#### Dimensionality estimation

To estimate the dimensionality of the latent manifold, we used two methods. To first methods leverages the geometric properties of the geodesic distance matrix ρ(i, j). We specifically studied the statistics of nearest neighbour distances (cf. Fig. 2C and Fig. 3C). Suppose that the neural states were confined to a two-dimensional sheet in high-dimensional neural state space. Within the sheet, the cumulative number of points N within distance r will increase quadratically with distance r, as more points on the sheet will fall within the neighbourhood, thus recovering the two-dimensional sheet structure. Using this variation of the correlation dimension, we found a wide range of values for which the number of points scaled like a power law. We fit this power law by minimizing the quadratic error to the model function N(r) = c r^d^, in which N is the total number of neighbours, r is the distance, and c and d are fit parameters. The second method uses the reconstructed held-out data. For every held-out trial, we obtained a curve of R^2^ vs the embedding dimension (cf. Fig. 2E and Fig. 3E - left panel). We fit to this curve a step-wise linear function that increased linearly up to a point, and was flat beyond. Fitting this model function to every held-out trial produces a large number of estimates for the “knee” beyond which increasing the embedding dimension led to negligible improvements in reconstruction. Finally, we estimated the mean and C.I. of these values using bootstrapping (cf. Fig. 2E, 3E - right panel).

#### Decoding of behavioral variables

We used Gaussian Process Regression (GPR) as the primary nonlinear regression technique to learn a function from latent space to position and evidence. Other nonlinear regression methods such as LLE yielded similar results, whereas linear decoding methods generally failed. Figures 2J,K and 3J,K show the correlation coefficients between the position and evidence values in the behavioural session of each mouse predicted from the learned regression model and true position and evidence values (averaged across the 10 folds of 10-fold cross validation). Shuffles were computed by random circular shifts of the test-fold. Only for visualization, Figures 2G,H,I and 3G,H,I were smoothed across the 10 nearest neighbours in latent space. No smoothing was applied to the example manifold firing fields in Figure 2F and 3F.

#### Hyperalignment analyses

To align neural data (Fig. 4) across animals and brain regions, we analyzed manifolds obtained from entire sessions (for an analysis timepoint-by-timepoint, see next paragraph). Hyperalignment across two mice was performed as follows: We first fit the neural data of mouse A with MIND to obtain a set of T d-dimensional latents x_A_(t). We then perform GPR to learn a map from the d-dimensional latents to a behavioural variable e_A_(t) = GPR[x_A_(t)]. Next, we perform MIND on the data of mouse B. This yields a different set of d-dimensional latent vectors x_B_(t). From these latents, we predict the behaviour of mouse B using the GPR trained on mouse A and a rotation matrix R with e_B_(t) = GPR[R x_B_(t)]. We performed this comparison based on the five-dimensional embedding, which made the mPFC and HPC manifolds comparable. The rotation matrix R was calculated from a five-dimensional representation of the SO(5) so that R = Π_i_ expm(g_i_ c_i_). The function expm() is the matrix-exponential of g_i_, the ten generators of the SO(5), multiplied with a scalar angular parameter c_i_. The c_i_ are the ten rotation angles that are optimized for optimal alignment. These rotations were cross-validated by optimizing the angles c_i_ on the first half of the data and decoding of position and evidence on the second half using the c_i_ from the first half. The dataset contained 10 recordings of mPFC and 8 recordings of HPC. For every single recording, we decoded position and evidence using the hyperaligned methods described above based on the other datasets. The results were then organized into three categories: The best decoding from the same animal (left column in Fig 4B,C), the same brain region but different animal (middle column in Fig 4B,C), and a different brain region across all animal (right column in Fig 4B,C). The thin black lines are individual datasets; eight for HPC and ten for mPFC. The results were statistically indistinguishable.

#### Across-region decoding and timing analyses

For the examples in Fig. 5A,B,C and Fig. 5 H,I,J, we plotted the respective latents as a heatmap. Consistent with the examples in Fig. 2 and 3, these heatmaps were smoothed across the ten nearest neighbors for visualization purposes. Neither the following example firing fields, nor the following analyses were smoothed.

To decode latent variables across areas on a per-timepoint basis, we followed the same method that we used to decode position and evidence from the latent: 10-fold cross-validated Gaussian Process Regression. To predict the PFC latents from HPC, we separately predicted the six latents of the six-dimensional embedding of mPFC, and computed a correlation coefficient between the held-out data and the prediction. The results were then just averaged across the 10 held-out folds. Small gray dots in Fig. 5F show the averages of the best mPFC latent decoding for each individual dataset. Blue bars are the mean across datasets. Errorbars show the standard deviation across datasets. The analysis for Fig. 5M was similar, but we predicted the five latents of the five-dimensional HPC manifolds.

For timing analysis, the same method as above was used, but the two time series of mPFC and HPC latents were shifted relative to each other. In Fig. 5G, we predicted the shifted six-dimensional embedding of mPFC from the five-dimensional embedding of HPC. For Fig. 5N we predicted the shifted five-dimensional embedding of the HPC manifold based on the six-dimensional embedding of the mPFC manifold.

To determine the fraction of across-area alignment caused by tuned neural responses to evidence and position (Suppl. Fig. 5), we computed residual trajectory (“R-Trajectory” in the following), based on the difference of the observed latent trajectory, and the prediction based on the experimentally controlled variables like position and evidence. In other words, the R-Trajectory captures neural dynamic above and beyond tuning to these physical and learned variables. This was done in a cross-validated way. We split the dataset into ten blocks, learned a GPR regression model based on position and evidence on nine training blocks, and predicted the manifold coordinates in the test block. All ten test blocks were then concatenated to obtain the cross-validated expected latent trajectory based on behavior. We then subtracted this control manifold from the data, and repeated the across-area decoding analyses (see above) for different embedding dimensions (Suppl. Fig. 5B,D). To compare these results with the full trajectories (Fig. 5F,M), we computed the ratio and found that around 50% of the observed manifold alignment is caused by tuning to position and evidence.

Finally we tested for timing differences in subphases of the task (Suppl. Fig. 6). To this end, we repeated the timing analyses in the Cue Period, defined as the part of the maze between 0cm and 200 cm, and the delay region, defined as 200 cm and 300 cm.

#### Statistical Tests

All statistical tests were performed with MATLAB (2015b, 2018a, 2018b and 2020a; Mathworks). Bonferroni correction of P values was performed by multiplying the unadjusted P value by the number of multiple comparisons made. In cases in which the corrected P value exceeded 1.0, we reported the value as 1.0.

## Supplemental Figures

**Suppl. Fig. 1:**
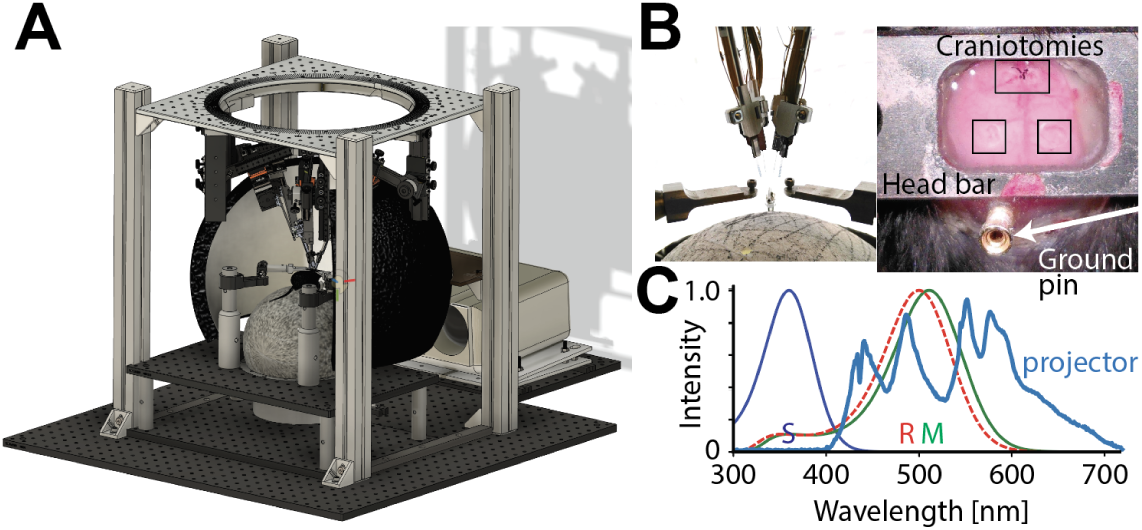
Setup for acute recordings in Virtual Reality. **A)** CAD drawing of the setup. Note the micro-manipulators mounted about the mouse, the styrofoam projection dome around the animal, and the VR projector. **B)** Left: Photograph showing arrangement of probe holders and head bars. Note the typical angles of insertion. Right: Microscope image of the craniotomies and the ground pin. **C)** VR light spectrum compared to mouse short and middle wavelength cone and rod fundamentals^40^.

**Suppl. Fig. 2:**
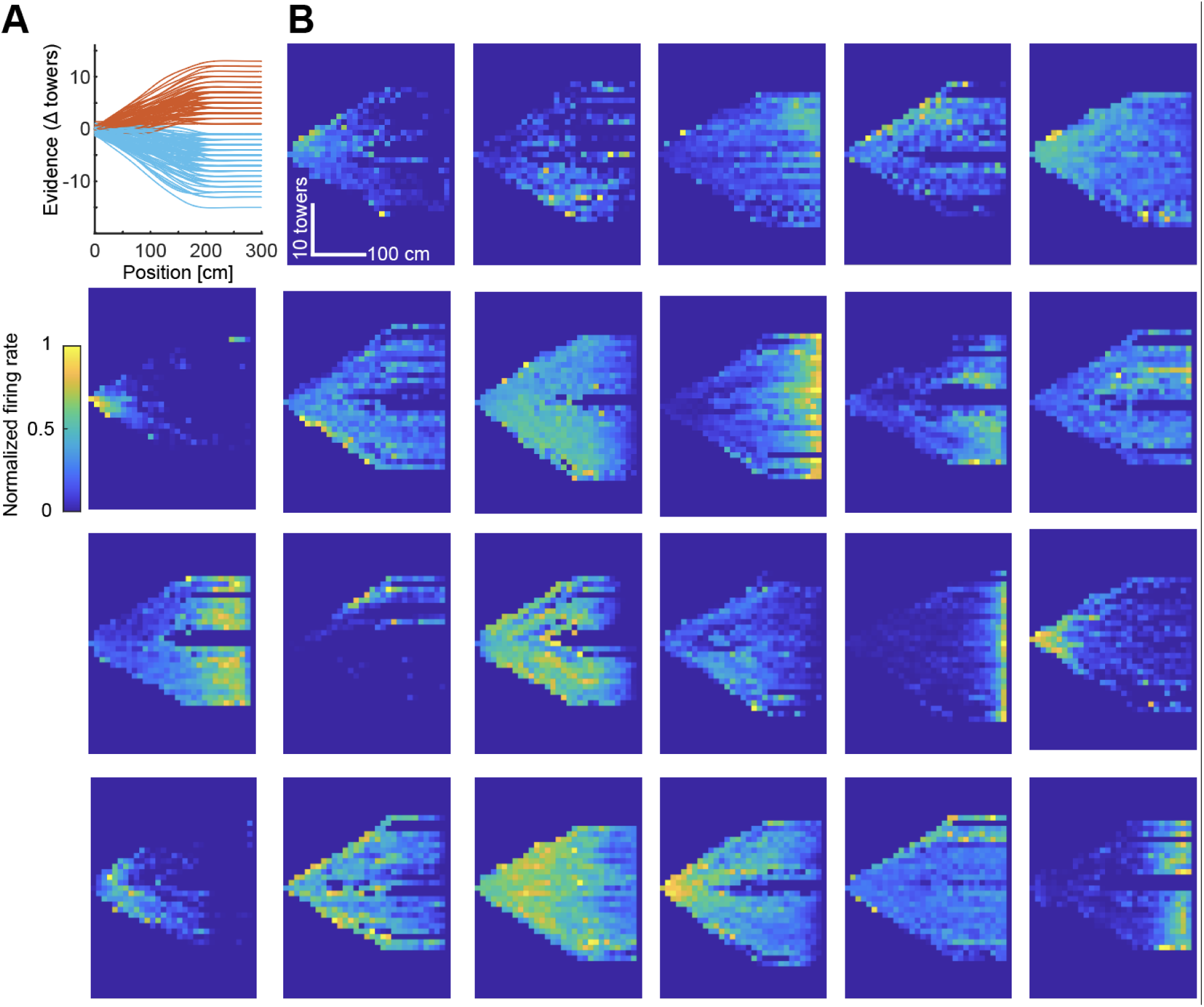
Example tuning curves in mPFC. **A)** Trials in position/evidence space constitute trajectories (thin lines), and indicate the correct decision (red/blue). Note the triangular shape imposed by the task structure: Large evidence values cannot be encountered early in the maze. **B)** Binned firing of 23 example neurons in position/evidence space.

**Suppl. Fig. 3:**
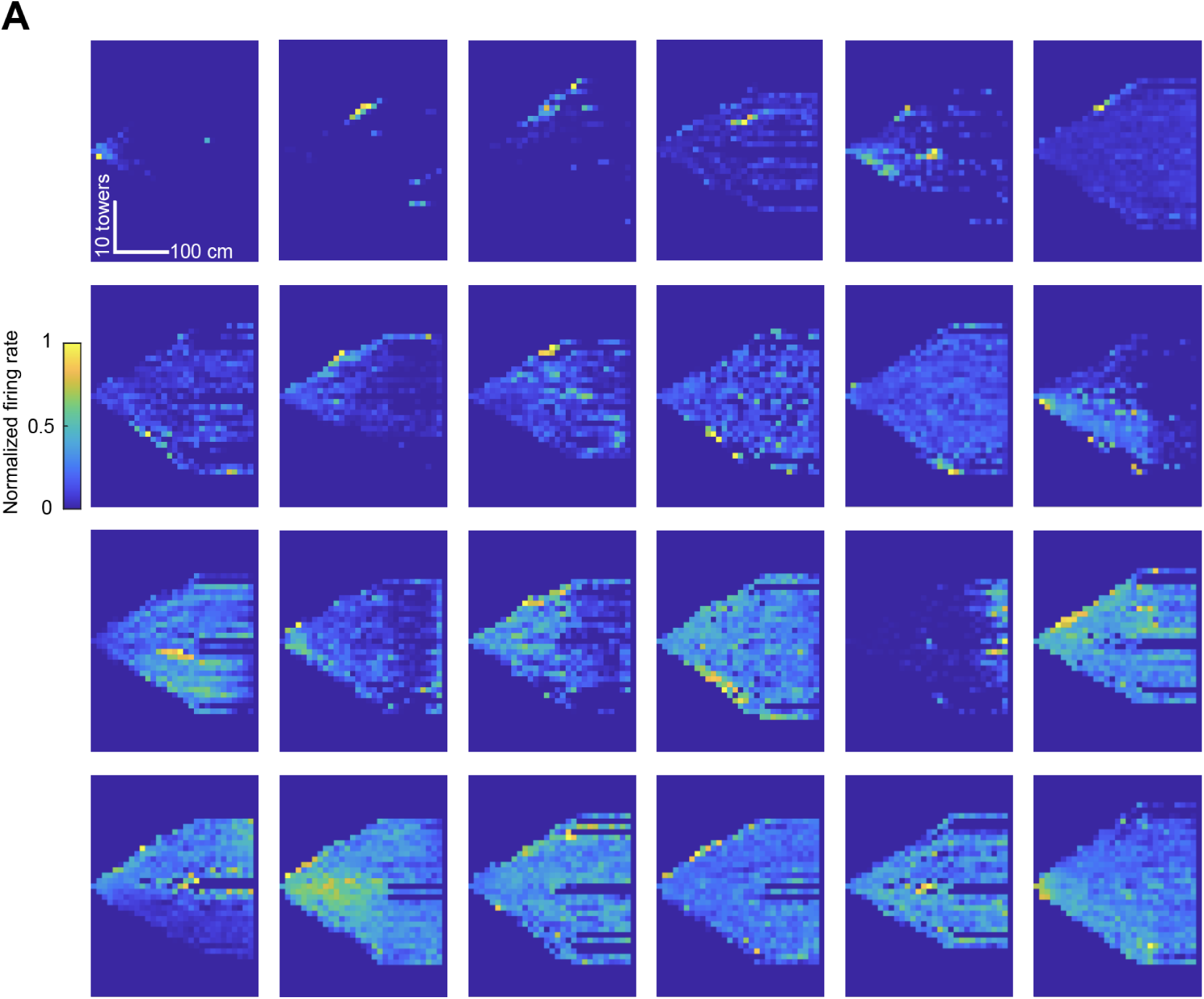
Example tuning curves in HPC. **A)** Binned firing of 24 example hippocampal neurons in position/evidence space. Note the tight firing fields in comparison to the mPFC data. An analysis of tuning properties is provided in^11^.

**Suppl. Fig. 4:**
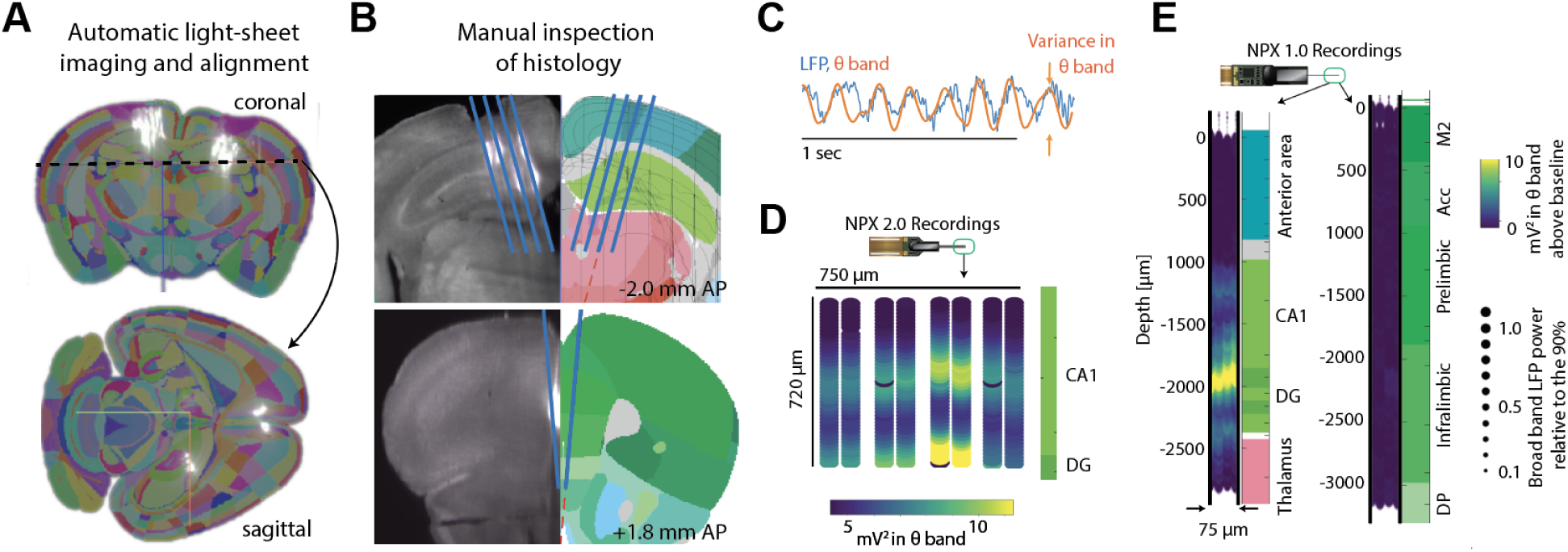
Probe Alignment procedure. **A)** Light-sheet image of an example brain. Shown is fluorescences with overlaid Allen Atlas morphed to a best fit. The coronal and sagittal sections illustrate the probe locations in Hippocampus and medial frontal cortex. **B)** Manual inspection to validate probe placement. **C)** To precisely align probe location to a reference atlas, we bandpass filtered all LFP data in the theta band (6-10 Hz) and measured the power in the theta band. **D)** Example of theta power maps across the four shanks of a 2.0 probe inserted into Hippocampus. Two bands are visible corresponding to CA1, and the dorsal part of the Dentate Gyrus. Note the missing dot. This corresponds to Channel 127, which is not a recording channel regardless of referencing. The rest of the electrodes were specified through a shank map that selected the bottom 96 electrodes across the four shanks. **E)** Example for two recordings made with Neuropixels type 1.0. The left recording was a probe inserted through Hippocampus. The right recording shows mPFC. Highest power in the theta band was found around the str. lacunosum-moleculare of the hippocampal CA1, located just above the Dentate Gyrus.

**Suppl. Fig. 5:**
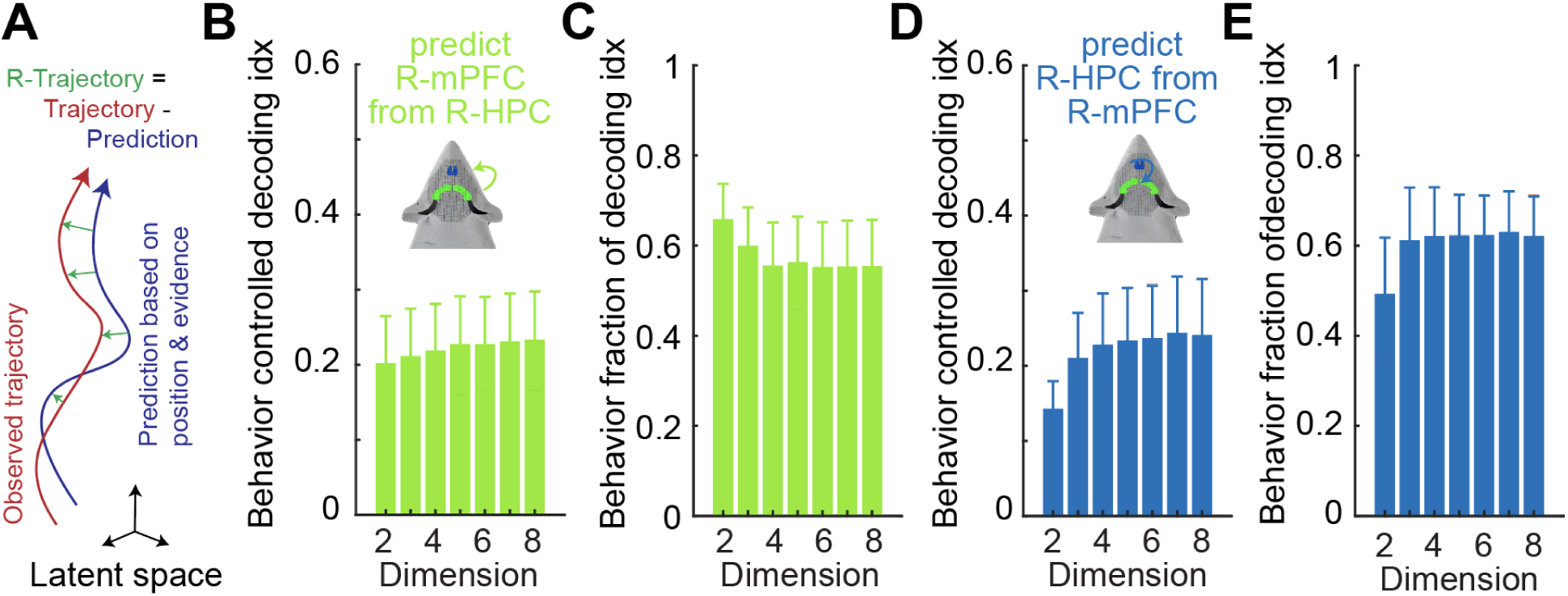
Determining the fraction of across-area alignment of the manifolds caused by shared tuning to position and evidence. **A)** Cartoon of how residual trajectories (“R-Trajectories”) are computed, based on the difference of the observed latent trajectory, and the prediction based on the experimentally controlled variables position and evidence. **B)** Running the across-area decoding analysis on R-trajectories for different embedding dimensions, predicting the residual mPFC latents from the residual HPC latents. **C)** The ratio of B to the full trajectories (cf. Fig. 5F,M). Note that around 50% of the observed manifold alignment is caused by tuning to position and evidence. **D)** Same as B but predicting HPC from mPFC for various embedding dimensions. **E)** Same fraction explained as C, but for the HPC from mPFC prediction.

**Suppl. Fig. 6:**
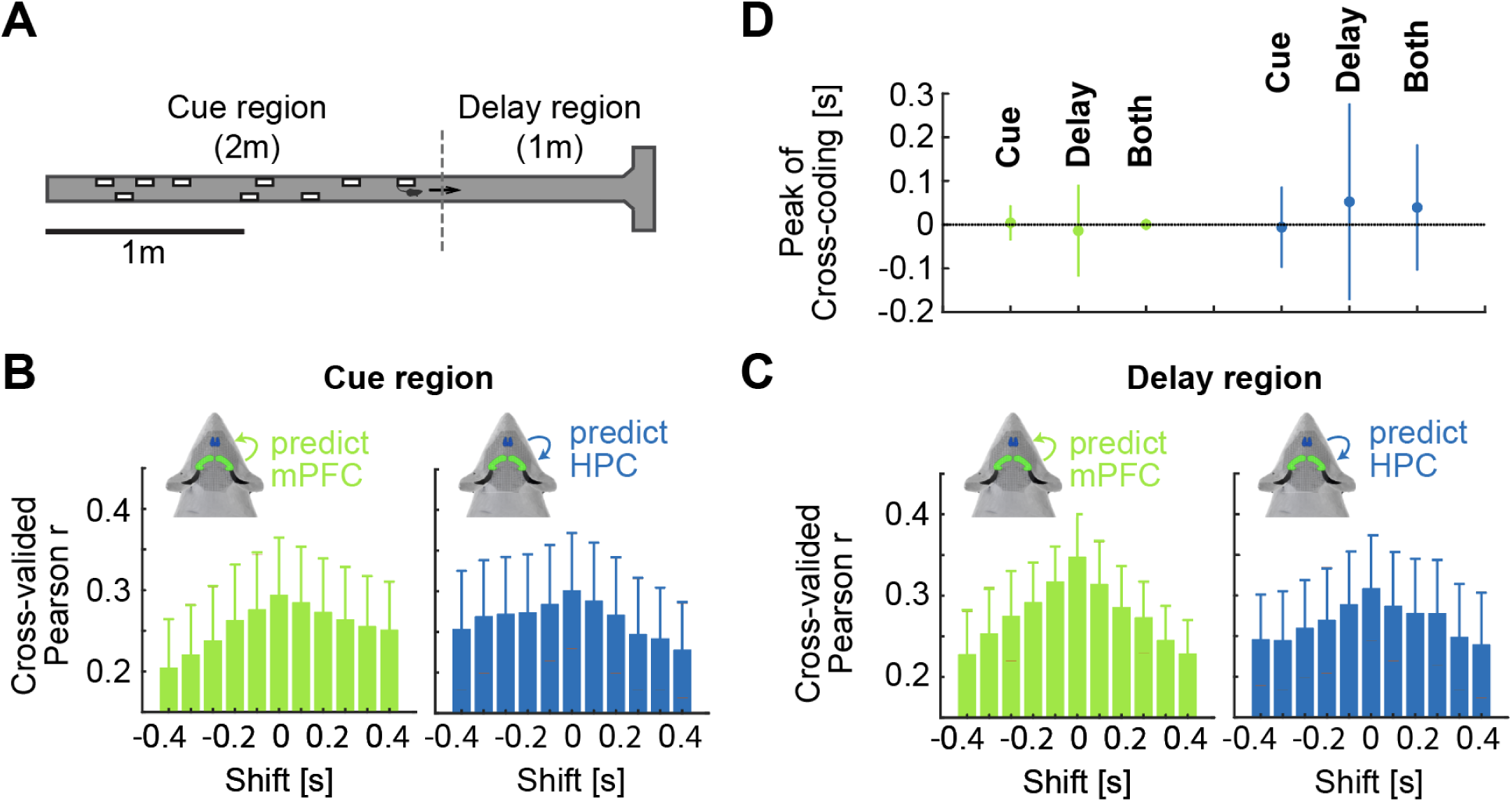
Delay measurements split by Cue and Delay period. **A)** Cartoon of the accumulating towers task, illustrating the phases in which the cues are presented (first two meters), and the region in which the mouse has to remember the correct side (another meter). **B)** Decoding of the mPFC latents from the HPC manifold (left; green) and vice versa (right; blue) for different time shifts, but only in the Cue period. Notice the largest peak at shift zero in both directions. The same analysis for all data is shown in Fig. 5G and Fig. 5N. **C**) Same as B, but during the Delay period. **D)** Comparison of the optimal timing delay of the cross-area decoding in the two phases vs. all data. Shown are means and S.E.M. Notice the larger error bars, caused by splitting the data into Cue and Delay sections. Green is the optimal timing delay predicting mPFC based on HPC, and blue vice versa. Both are statistically indistinguishable from the absence of a timing delay.

